# Antimicrobial resistant *Salmonella* spp. circulating in antibiotic-free organic pig farms of northern-Thailand

**DOI:** 10.1101/2020.12.16.419408

**Authors:** Pakpoom Tadee, Prapas Patchanee, Ben Pascoe, Samuel K. Sheppard, Dethaloun Meunsene, Phacharaporn Tadee

## Abstract

We investigated the prevalence of *Salmonella* circulating in local organic pig farms in northern Thailand and typed isolated clones to better understand the population structure of the underlying *Salmonella* contamination. In total, 112 samples from 11 organic pig farms were processed from October to December 2018. *Salmonella* were detected in 9 farms with an overall prevalence of 25.0% (28/112). Prevalence detected in fecal, feeder, and boot swab samples was found to be 32.7% (17/53), 17.7% (6/34), and 20.0% (5/25), respectively. Of the 28 positive strains, Seven *Salmonella* serotypes were identified, with *S*. Rissen being the most common (15/28; 53.6%). 89.3% (25/28), 78.6% (22/28) and 71.4% (20/28) of isolated *Salmonella* were resisted against tetracycline, ampicillin, and sulfamethoxazole-trimethoprim, respectively. From multilocus sequence typing (MLST) analysis, the phylogenetic tree suggests dissemination of specific clones within herds that share routes of pig transportation and point mutations in housekeeping genes within herds. A minimum spanning tree (MST) revealed that *Salmonella* contamination in organic pig farming is possibly linked with conventional farming. Based on the own results, strictly highly organic practices provide a safe alternative enhancing domestic consumer trust and improve public health safety.

## 1. Introduction

Foodborne illness caused by non-typhoidal *Salmonella* is an important public health issue and is a significant economic burden worldwide [1-5]. Transmission of the pathogen to humans occurs mainly through contaminated food products from livestock [6,7], including consumption of pig or pork products [8,9]. Clinical infection manifests with the onset of fever, followed by gastrointestinal tract disorders such as nausea, vomiting, and profuse diarrhea within 8–48 hours [10]. Multidrug-resistant *Salmonella* can complicate treatment and is a serious rising issue in public health [11], leading to a reduction in first line empirical treatment efficacy, limiting treatment choices, and prolonging illness [5].

*Salmonella* has been identified at all stages of the food production chain [9]. Initial contamination pressure at farm level is directly related to continued contamination at subsequent levels, through slaughtering, retail, and the final cooking process [12,13]. Pigs are often cited as the primary source of human infection, as *Salmonella* can multiply in the pig’s gut and spread between animals via the fecal–oral route directly or through the environment. Healthy pigs can also carry *Salmonella* without showing any signs of infection [14]; thus, increasing the opportunity for spread of *Salmonella* into the human food chain.

Currently, increasing demand has led to pig production being transformed into an intensified industry. Large numbers of antimicrobials have been implemented [15] and widespread usage and misuse have contributed to the emergence of drug-resistant bacteria [16], one of several concerns that has fueled the increase in “organic” farming. Pigs are produced under natural management, where antibiotics and synthetic hormones are prohibited. It has set itself the goal of instituting high-quality yields with environmentally friendly production under animal welfare standards [17,18]. In northern Thailand, local organic pig farms have been established for half a decade. However, data on the *Salmonella* contamination or any resistance situations have not been clearly determined.

Bacterial typing is essential for effective disease investigation and surveillance [19]. Serotyping is based on immune reaction raised in the host and has limited discriminatory power to distinguish *Salmonella* clones [20,21]. Multilocus sequence typing is able to overcome limitations in detection and help distinguish between *Salmonella* clones and aid outbreak assessment. The method is also more portable and by using sequence data that can be compared between laboratories, strains can be characterized nationally or internationally [22,23]. Large global studies use this method for epidemiological studies that clearly and precisely differentiate bacterial genotypic diversity in different hosts and geographical distributions [21,24].

The aims of this study were to determine the prevalence of *Salmonella* spp. circulating in organic pig farms in northern Thailand, and evaluate diversity of the pathogen. Phenotypic characteristics (serotypes and antimicrobial resistance patterns) and genotypic sequence types were measured, to understand the local population structure and dynamic propagation. The results will enhance domestic consumer trust and help improve public health safety in the region.

## 2. Material and Methods

### 2.1. Sample collection

This study was conducted under ethical approval reference number R13/2561 from the Animal Care and Use Committee of the Faculty of Veterinary Medicine, Chiang Mai University (FVM-ACUC). Pig fecal samples, feeder swabs and workers’ boot swabs were collected from local organic pig farms in Chiang Mai, Chiang Rai, and Lamphun provinces, Thailand from October to December 2018. To estimate the sample size in a single proportion, the sample size of this study was determined using the Win Epi online program at (http://www.winepi.net/uk/index.htm) [25]. Because of entering 50% for the estimated prevalence will result in the highest sample size. Therefore, a rate of 50% was used as the “expected prevalence (*p*)”. Ten percent and 0.95 were selected as the “desired precision (*E*)” and “confidence level (*Z*_1−*α*/2_)” values, respectively. For the infinite population, at least 97 samples are designated. However, at the end, 112 samples were carefully chosen to provide greater accuracy and reliability of the investigation results.

Eleven open housing system pig farms with a herd size range of 30–60 pigs were randomly selected in the study. The selection criteria of all organic farms were: (1) no antibiotic or hormone usage throughout the pigs lifespan: (2) all piglets were born under the organic system; (3) use of natural feed ingredients from an organic agriculture system (either with or without commercial feed); (4) farmed according to animal welfare standards; and (5) either natural mating or artificial insemination for breeding. Approximately ten samples (5 pig feces from digital rectal examination, 3 feeder swabs, and 2 workers’ boot swabs) were collected from each farm. For fecal and feeder swab samples, up to two samples/pen were chosen. All collected samples were kept in separate sterile bags and transferred to the laboratory under chilled conditions.

### 2.2. Salmonella isolation and identification

Isolation and identification of *Salmonella* spp. were performed following the ISO 6579:2002 Amendment 1:2007, Annex D technique, the detection of *Salmonella* spp. in animal feces, and environmental samples from the primary production stage [26]. Fresh fecal samples were diluted with nine times buffered peptone water (BPW) (Merck, Germany) and 25 g of sample was added to 225 mL of BPW pre-enrichment media. Swab samples were diluted with 100 mL of BPW. Mixtures were homogenized for 2 min and an aliquot of 0.1 mL was dropped on Modified Semi-solid Rappaport-Vassiliadis (MSRV) (Oxiod, United Kingdom) plates and incubated at 42 °C for 24 h. The culture material surrounded by a turbid ring was then streaked on Xylose Lysine Deoxycholate (XLD) agar (Oxiod, United Kingdom), and Brilliant Green Phenol Red Lactose Saccharose (BPLS) agar (Merck, Germany) with incubation at 37 °C for 24 h. Presumptive *Salmonella* colonies were additionally checked by biochemical measurement, including testing of triple sugar iron (TSI) (Oxiod, United Kingdom), urease, and motile indole lysine (MIL) decarboxylase (Merck, Germany). *Salmonella* positive samples were recorded. Only one isolate from each sample was chosen for further investigation.

### 2.3. Serotyping and Antimicrobial susceptibility testing

All detected *Salmonella* spp. were serotyped by serum-agglutination according to the White–Kauffmann–Le Minor scheme [20]. Antimicrobial susceptibility testing was determined by agar disk diffusion techniques with ten panels of antimicrobial agents, comprising amoxicillin-clavulanic acid 20/10 μg, ampicillin (AMP) 10 μg, chloramphenicol (C) 30 μg, ciprofloxacin (CIP) 5 μg, cefotaxime (CTX) 30 μg, nalidixic acid (NA) 30 μg, norfloxacin (NOR) 10 μg, streptomycin (S) 10 μg, sulfamethoxazole-trimethoprim (SXT) 23.75/1.25 μg, and tetracycline (TE) 30 μg [27]. Strains with intermediate resistance profiles were recorded as resistant strains to avoid underestimation.

### 2.4. Statistical analysis

*Salmonella* prevalence with their 95% confidence levels were determined by descriptive statistics. Comparisons between groups of sample types were performed using Fisher’s exact test. All analyses were accomplished using Epi Info™, version 7 (Centers for Disease Control and Prevention, USA). Statistical significance levels were measured at p < 0.05.

### 2.5. Multilocus sequence typing

All *Salmonella* strains detected were genotyped into sequence types (STs) by the MLST technique. Total genomic DNA was extracted according the protocol described by Liu *et al*. (2011) [21]. Seven housekeeping genes, including *aro*C (chorismate synthase), *dna*N (DNA polymerase III beta subunit), *hem*D (uroporphyrinogenIII cosynthase), *pur*E (phosphoribosylaminoimidazole carboxylase), *suc*A (alpha ketoglutarate dehydrogenase), *his*D (histidinol dehydrogenase), and *thr*A (aspartokinase I/homoserine dehydrogenase) were taken for MLST profiling. Polymerase Chain Reaction (PCR) amplification of all seven genes was accomplished using the methods previously described by Achtman *et al*. (2012) [28]. Then, the PCR products were sent for sequencing to the Macrogen Service Center, Republic of Korea. All sequences obtained in each gene were transformed into allele number and compiled for sequence type (ST) data from the database of http://mlst.warwick.ac.uk/mlst/ [29].

Using Bionumerics^®^ software, version 7.6 (Applied Maths, Belgium), by the unweighted pair group method with arithmetic mean algorithms (UPGMA) through individual similarity matrices, cluster analyses of the seven genes were executed. Local epidemiological results of strains currently circulating in organic pig farms in the study area were displayed in phylogenetic networks. In addition, to develop understanding of evolution and population structure, minimum spanning tree (MST) analysis was used to analyze the genetic relationship between *Salmonella* stains obtained from this study together with the geographically and temporally matched strains previously submitted to the MLST database [29], Warwick Medical School, University of Warwick. Those included were Thai strains recovered from pig (*n* = 218), human (*n* = 42), plant (*n* = 37), frozen food (*n* = 30), aquatic animal (*n* = 16), dried food (*n* = 13), poultry (*n* = 9), soil (*n* = 6), invertebrate (*n* = 2), cooked food (*n* = 1), and marine mammal (*n* = 1), submitted during 2009–2018. The STs, which are closely associated in loci characteristics, were demonstrated close together.

## Results

### 3.1. Salmonella prevalence

From October to December 2018, 112 fecal and environmental samples were collected from 11 local organic pig farms (farms A–K). Eight to 13 samples were obtained from each. The overall prevalence of *Salmonella* identified in northern Thailand was 25.00% (28/112; 95% CI: 17.30–34.07%). The prevalence of fecal, feeder swabs, and boot swabs were found to be 32.08% (17/53; 95% CI: 19.92–46.32%), 17.65% (6/34; 95% CI: 6.76–34.53%), and 20.00% (5/25; 95% CI: 6.83–40.70%), respectively. There was no association between *Salmonella* frequencies detected and types of sample taken. *Salmonella* contamination was detected in nine of the eleven farms tested, with only two farms (farms F and K) negative. Between 10–20% of the samples obtained from six farms (farms B, D, E, G, H, and I) were positive to *Salmonella*. For the three remaining farms (farms A, C, and J), 50–60% detection rates were distributed.

### 3.2. Salmonella sero-distribution

Seven *Salmonella* serotypes were identified in the study. *S*. Rissen was the most common (15/28; 53.57%), followed by monophasic *S*. Typhimurium 1,4,[5],12: i: – (7/28; 25%), *S*. Weltevreden (2/28; 7.14%), and one strain (1/28; 3.57%) each of *S*. Hvittingfos, *S*. Krefeld, *S*. Stanley, and *S*. Typhimurium.

### 3.3. Antimicrobial susceptibility testing

Of all 28 strains taken from *Salmonella* positive samples, only one (3.57%) was susceptible to all antimicrobials tested, while 24 strains (85.71%) were resistant to at least three different antimicrobials, classifying them as multidrug-resistant (MDR) strains. Resistance of *Salmonella* from the current study could be found concerning eight of ten antimicrobials tested. No resistance was observed to ciprofloxacin and norfloxacin. Most strains were resistant to tetracycline (25 strains; 89.29%), followed by ampicillin (22 strains; 78.57%) and sulfamethoxazole-trimethoprim (20 strains; 71.43%) (**Figure 1**).

**Figure 1.**
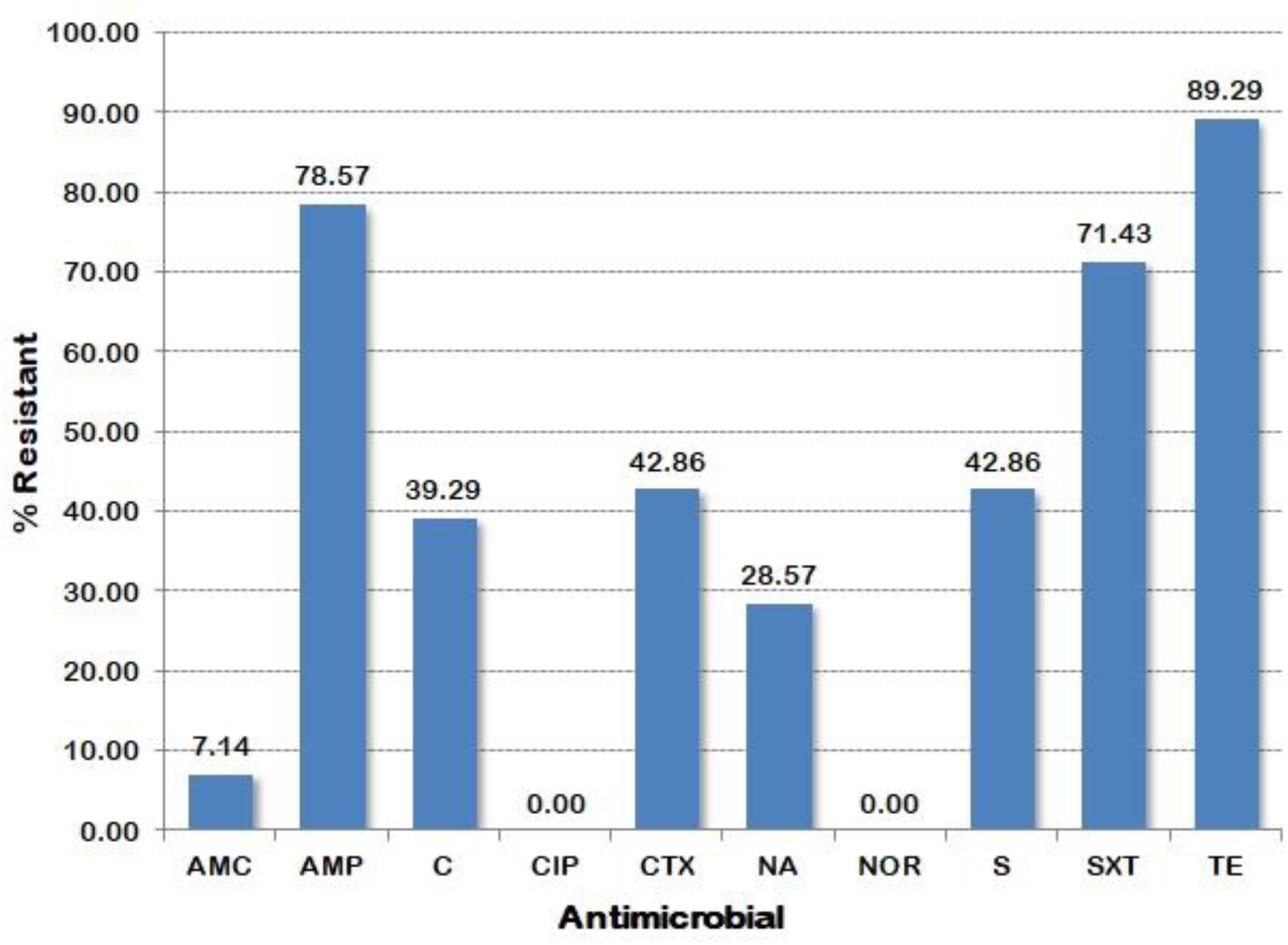
Antimicrobial resistance rates of *Salmonella* strains isolated from local organic pig farming in northern Thailand. Antibiotic abbreviations: amoxicillin-clavulanic acid (AMC); ampicillin (AMP); chloramphenicol (C); ciprofloxacin (CIP); cefotaxime (CTX); nalidixic acid (NA); norfloxacin (NOR); sulfamethoxazole-trimethoprim (SXT); streptomycin (S); tetracycline (TE)

### 3.4. Multilocus sequence typing

Sequencing data of seven housekeeping genes of all strains were submitted to the *Salmonella* spp. definition database to query allelic numbers and corresponding STs. In total, 12 different genotypic characters were assigned. Six of them were matched with the previously described in MLST database [29], could be assigned in ST number. The majority were of ST469 (*n* = 14), followed by three strains of ST34 and two strains of ST365. Six others could not be assigned a ST, highlighting the diversity of *Salmonella* isolates found on these farms. Variation in the *suc*A and/or *his*D or *thr*A loci enabled assignment to ST complex or clonal complex (CC) level. Five strains were assigned to CC1, and the other one was CC66. The genetic relationship between all 28 *Salmonella* strains was explored using sequence variation from the seven housekeeping genes with UPGMA algorithms (**Figure 2**).

**Figure 2.**
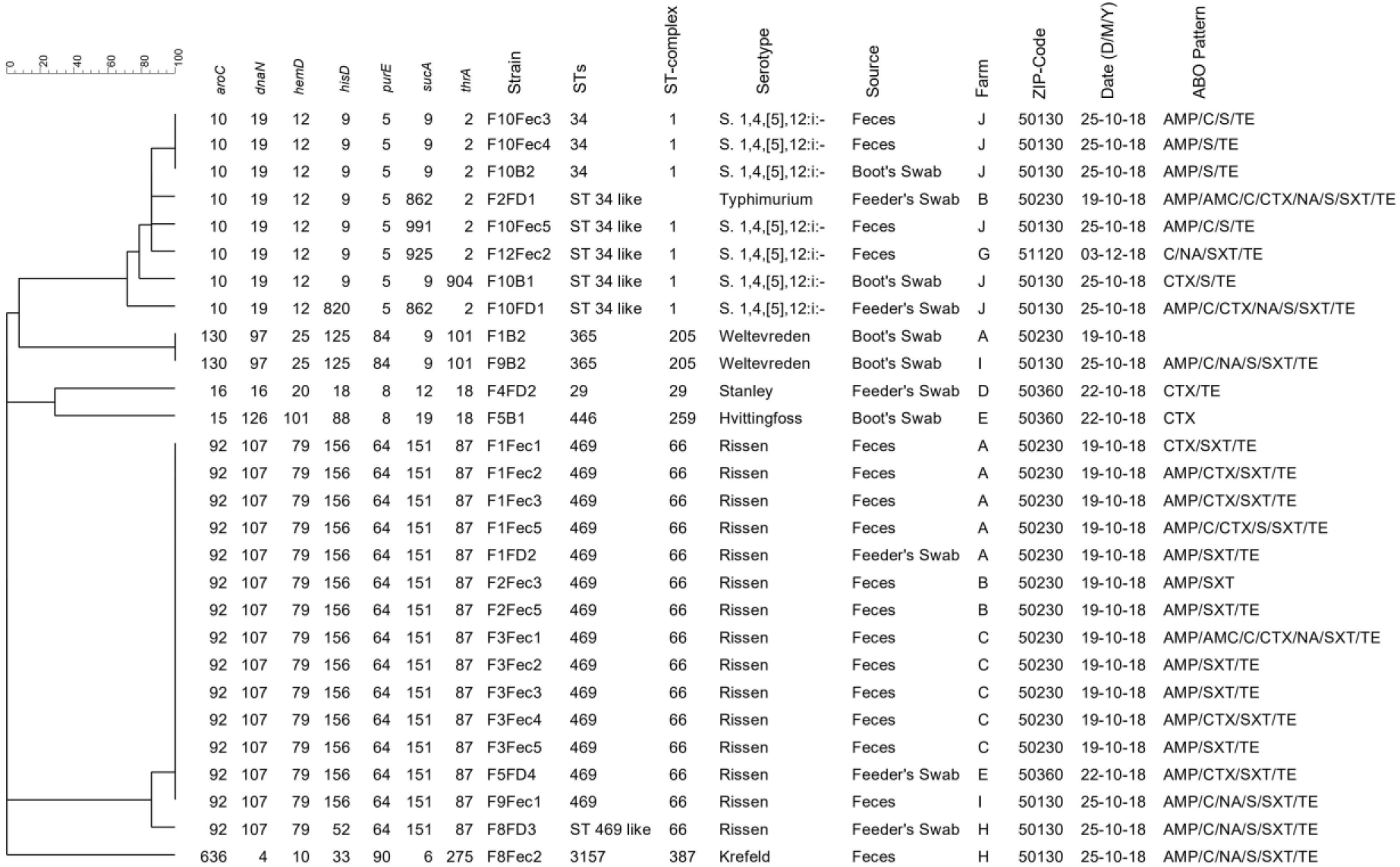
Dendrograms generated using UPGMA algorithms based on MLST profiles with the phenotypic characterization and epidemiological data of *Salmonella* circulating on organic pig farms in northern Thailand.

Six clusters were determined with two or more MSLT gene differences, representing different CCs. The strains grouped in the dominant serovar Rissen ST469 were mostly obtained from three nearby farms (farms A–C, which each has a similar zip-code), and two identical strains were also obtained from farms approximately 30–70 km away from each other (farms E and I). Besides, strains grouped in Weltevrenden ST365 were also obtained from farms spread more than 30 km apart (farms A and I).

Serovar Rissen ST469 from farm A was recovered from feeder swabs and fecal samples. Similar to the ST34 strains from farm J, the strains were recovered from a boot swab and fecal samples. Moreover, in farms A, B, E, H, I, and J, at least two ST characteristics were identified.

In order to better understand the epidemiology of the pathogen, a minimum spanning tree (MST) was constructed to analyze the 28 strains alongside 375 Thai strains previously submitted to the MLST database in the ten-year period, 2009–2018 (**Figure 3**). A total of 403 Thai strains was constructed describing a population from 11 different hosts. Seventy-one STs were distributed. More than two-fifths of the strains were closely related and shared allelic profiles in at least five loci, with similar CCs shaded in gray. ST34 was the most frequently detected, followed by ST465, ST365, and ST29. Members grouped in the top four STs were recovered from more than one host origin, such as organic pig, pig, plant, soil, poultry, food, and human. For the members grouped in ST365, one was obtained from an invertebrate. On the other hand, some STs were unique to a single host origin; ST185, ST410, ST590 were specific to plant sources. Similarly, ST1 and ST292 were specific to human and aquatic animals, respectively.

**Figure 3.**
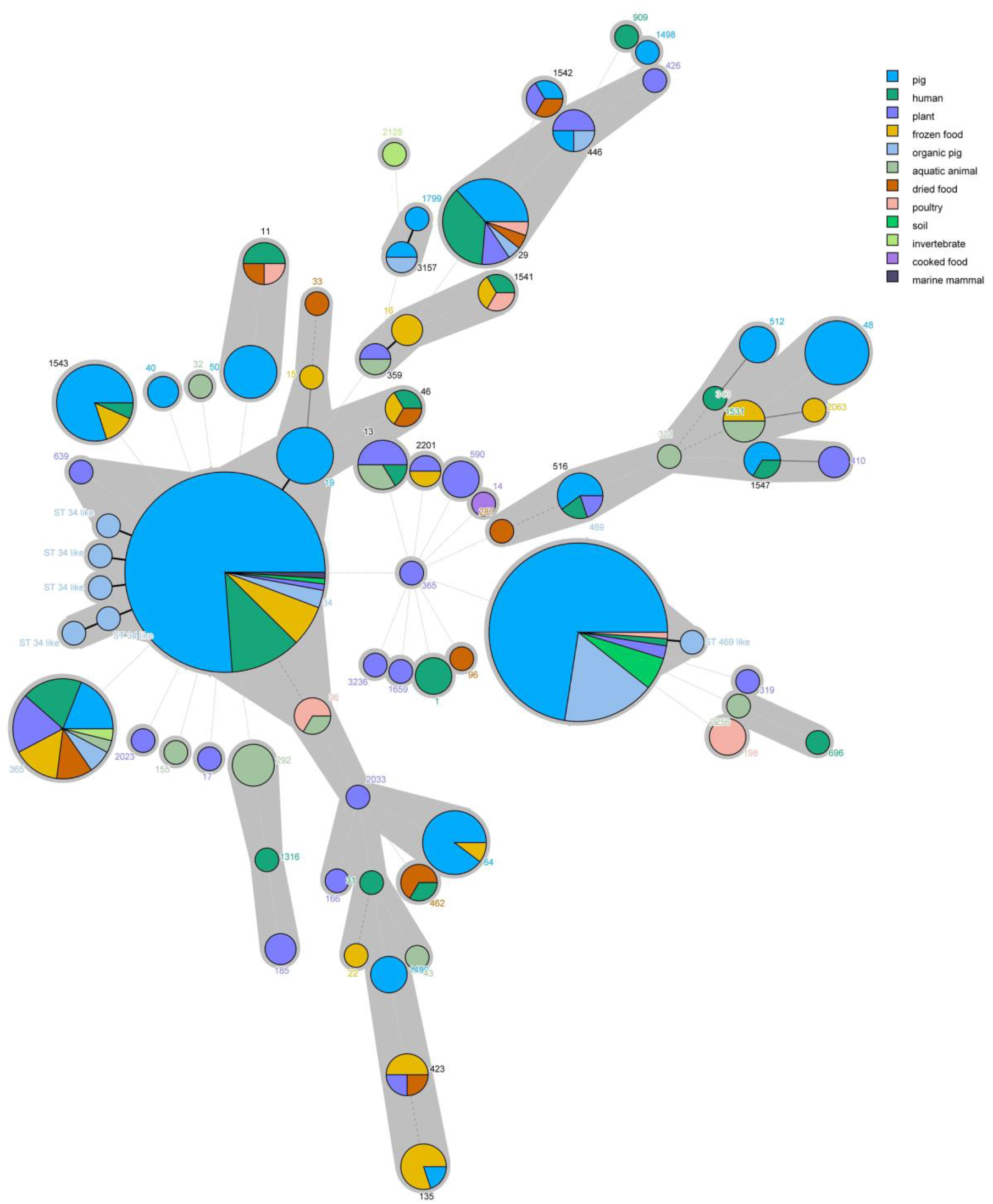
Minimum spanning tree (MST) analysis of 403 Thai population *Salmonella* strains recovered from various host origins during 2009–2018. Size of circles indicates the number of strains grouping in the same STs. Branch styles coding: thick black solid lines connect ST types with one locus difference; thin black solid lines connect ST types with double and three loci differences; gray dashed lines connect four loci differences; gray dotted lines connect more than four loci differences, respectively. Gray shaded areas indicate at least five shared loci between STs (grouping in common ST complexes)

## Discussion

This is the first cross-sectional study of *Salmonella* spp. circulating in organic pig farming in the northern region of Thailand. We identify an overall prevalence of 25% (95% CI: 18–35%), which is only slightly lower than rates identified in conventional pig farms from the same region during the same time period [31], that reported a prevalence of 31% (95% CI: 27–33%). This is far higher than rates identified in other similar studies on organic farms from other countries in SE Asia, where a prevalence of 8% (95% CI: 6–10%) was reported in a study of Korean organic pig farms by Tamang *et al*. (2015) [30]. These high levels of *Salmonella* contamination suggest that there is a breakdown in organic practices in rural northern Thailand.

All samples acquired from farms F and K were negative. However, only ten samples were collected from each farm, so these farms cannot yet be considered *Salmonella*-free. Although all the farms followed organic principles, at the present time farm K is the only one in northern Thailand that has received official accreditation from the Thai government, suggesting that standards may be higher on these farms. Organic products may provide an alternative for health-conscious local consumers and one hundred percent use of natural feed ingredients is likely beneficial [12], with a direct affects on the pigs’ gut microbiota [32,33]. The use of low pH fermented grain and natural organic feed stuff are act as a prebiotic, promoting growth of lactic acid bacteria to the detriment of the preferred conditions for *Salmonella* [34,35].

As one might expect following comparison among sample types, *Salmonella* prevalence from fecal samples tended to be higher than the value acquired from boot and feeder swab samples and transmission is likely to be via the fecal–oral route [14]. Ingested *Salmonella* can be shed via feces within 2 h, without any clinical symptom and spread to the wider environment, such as stable floors or feeders [8,12]. Consequently, hygienic practices in stable cleaning, boot disinfecting, and feed management are very important in reducing the risk of contamination [32]. However, other on-farm stressor factors including handling for mating, transportation as well as high density in the rearing area are also considered as multifactorial effects that increase the chance of *Salmonella* shedding [33]. For that reason, effective herd management allowing natural behavior expression should be fulfilled. This could also be considered an explanation for why *Salmonella* could not be found on farms F and K, which seemingly had higher hygiene standards than the other farms, even though natural feeds were not 100% implemented on farm F.

All 28 detected *Salmonella* strains were susceptible to norfloxacin and ciprofloxacin, both of which are fluoroquinolones, which is necessary in parenteral treatment of severe gastroenteritis cases [36,37]. Fluoroquinolones remain the most effective antimicrobials for pig related salmonellosis in humans. However, nine-tenths of strains tested in the study were MDR. High resistance rates of antimicrobials recently used in livestock production, such as ampicillin, tetracycline, or sulfamethoxazole-trimethoprim were found. This is consistent with reports on *Salmonella* isolated from conventional pig farms in northern Thailand that also revealed high levels of resistance to these drugs [31,38]. Generally, antimicrobial usage is currently prohibited in all organic farms selected for this study. Therefore, on-farm antimicrobial use is not a main factor affecting bacterial resistance in this situation. This finding illustrates the complicated and widespread nature of resistance in the farm community [12]. Breeders reared on conventional farms that enter the organic system may already carry drug-resistant *Salmonella* in their guts, providing an opportunity for transmission and infection. Accordingly, it is important to note that a resistance has also emerged in antimicrobial-free herds [39]. In relation to farms F and K, breeders are produced on farm, thus limiting the possibility of pathogen transmission from off-farm sources.

Shared STs belonging to different sample types were identified, suggesting a common source of contamination. Within herds, cross transmission can occur via carriers or any contaminated materials and the probability of re-infection could occur continuously [12,32]. If we consider the geography of our collection, 12 strains of ST469 were recovered from three different farms (farms A, B, and C) located in nearby areas, with the same zip-code. Farm staff, environmental contamination, and the sharing of routes for pig transportation could play a significant role in dissemination. Nevertheless, associations of spatial relatedness were also noted at farms located far apart (farms E and I). As a previously noted, common supply chains such as breeders, gilts, or feed stuffs might be inferred as an important route. *S*. Rissen was the most common serotype detected and is a dominant clone *Salmonella* contamination of pig farms in northern Thailand over the last 15-years [1,31]. In view of this, *Salmonella* contamination of both conventional and organic pig farms is probably linked.

Diversity in MLST genotypes in some farms provides evidence that multiple strains are present within a single herd. Farms A, B, E, H, and I harbored at least two genotypical characteristics of *Salmonella*. This may represent different sources of infection. For another explanation, variation in one or two alleles can be explained by point mutations in the housekeeping genes during persistent colonization of the herd. This is potentially what we see with the ST34 strains from farm J, where only one to three nucleotide changes can be identified in the *suc*A gene. Using the allele sequence comparison analysis function of the pubMLST *Salmonella* spp. MLST database [40], point mutations can be specified: 155A>G of *thr*A alters *thr*A_2 to *thr*A_904, with 99.8% identity. With the same identity, a variation 127G>A of *sucA* shifts *suc*A_9 to *suc*A_862. Additionally, variations 15C>T, 18A>G, and 54C>T of *sucA* make over *suc*A_9 to *suc*A_925 with only 99.4% identity. This allelic profile has not yet been referenced in the database and a new ST profile will need to be added.

As part of expanding knowledge of regional epidemiology, results of the Thai MLST obtained from several host origins from ten collection years were displayed in MST. A diverse set of 403 *Salmonella* spp. strains were evaluated. Sixty percent of them were pig associated strains, supporting a strong interest in *Salmonella* contamination of pork products in the region. Strains grouped in the majority STs were obtained from hosts linked with the human food chain. Pig, chicken, and aquatic animals were the primary source of infection, which can then spread the pathogen to the environment [9,12]. In addition, most strains derived from organic pig farms were clonally related to the pig associated strains from the conventional farming system. Predominant types, especially ST34 and ST469 have been spread into pig farms across the country and have been a source of contamination in the pig population for over a decade.

One of the ST365 strains was recovered from invertebrate, which could be an additional potential reservoir of salmonellosis. Moreover, ST1 was found to be unique in the human host origin. Not surprisingly, all strains belonging to ST1 were derived from *Salmonella* Typhi, which is renowned as the host specific serotype for humans [22]. For STs which were unique to other origins, the low numbers of strains submitted does not reflect that these cannot be transmitted or spread to the others. This requires further study to develop a complete understanding of their epidemiology.

## 5. Conclusions

Consumers demand choice and healthier options. Organic products aim to provide increased food security, animal wellbeing as well as consideration of the effects on environmental sustainability. However, in this study *Salmonella* was detected in nine of the eleven farms tested. All farms tested claimed to follow organic principles, but only one had received official Thai government certification – from which no *Salmonella* contamination was detected. Although *Salmonella* prevalence was slightly lower in these organic farms, contamination with multidrug resistant clones suggests that domestic consumers should still be wary and that subsequent meat products still pose a public health risk.

## Supplementary Materials

Table S1: Frequency table of categorical explanatory details described in the study.

## Author Contributions

Conceptualization, Pakpoom Tadee and Prapas Patchanee; Formal analysis, Pakpoom Tadee and Phacharaporn Tadee; Funding acquisition, Pakpoom Tadee and Prapas Patchanee; Methodology, Pakpoom Tadee and Dethaloun Meunsene; Project administration, Phacharaporn Tadee; Visualization, Dethaloun Meunsene; Writing – original draft, Pakpoom Tadee and Phacharaporn Tadee; Writing – review & editing, Ben Pascoe and Samuel K. Sheppard.

## Acknowledgements

The organic farms and support of their staff are acknowledged. We would like to thank the Bioinformatics & Systems Biology Research Center, King Mongkut’s University of Technology Thonburi (KMUTT) for their help and advice, especially in determining *Salmonella* genotypes. Finally, we are very grateful to the faculty of Veterinary Medicine, Chiang Mai University that reach agreement to collaborate on this study.

